# The LUBAC participates in Lysophosphatidic Acid-induced NF-κB Activation

**DOI:** 10.1101/2020.02.13.948125

**Authors:** Tiphaine Douanne, Sarah Chapelier, Robert Rottapel, Julie Gavard, Nicolas Bidère

**Affiliations:** Université de Nantes, INSERM, CNRS, CRCINA, Team SOAP, F-440000 Nantes, France; Princess Margaret Cancer Centre, University Health Network, Toronto, Ontario, Canada; Institut de Cancérologie de l’Ouest, Site René Gauducheau, 44800 Saint-Herblain, France

## Abstract

The natural bioactive glycerophospholipid lysophosphatidic acid (LPA) binds to its cognate G protein-coupled receptors (GPCRs) on the cell surface to promote the activation of several transcription factors, including NF-κB. LPA-mediated activation of NF-κB relies on the formation of a signalosome that contains the scaffold CARMA3, the adaptor BCL10 and the paracaspase MALT1 (CBM complex). The CBM has been extensively studied in lymphocytes, where it links antigen receptors to NF-κB activation via the recruitment of the linear ubiquitin assembly complex (LUBAC), a tripartite complex of HOIP, HOIL1 and SHARPIN. Moreover, MALT1 cleaves the LUBAC subunit HOIL1 to further enhance NF-κB activation. However, the contribution of the LUBAC downstream of GPCRs has not been investigated. By using murine embryonic fibroblasts from mice deficient for HOIP, HOIL1 and SHARPIN, we report that the LUBAC is crucial for the activation of NF-κB in response to LPA. Further echoing the situation in lymphocytes, LPA unbridles the protease activity of MALT1, which cleaves HOIL1 at the Arginine 165. The expression of a MALT1-insensitive version of HOIL1 reveals that this processing is required for the optimal production of the NF-κB target cytokine interleukin-6. Lastly, we provide evidence that the guanine exchange factor GEF-H1 activated by LPA favors MALT1-mediated cleavage of HOIL1 and NF-κB signaling. Together, our results unveil a critical role for the LUBAC as a positive regulator of NF-κB signaling downstream of GPCRs.

## Introduction

Lysophosphatidic acid (LPA) is a natural serum phospholipid, which promotes growth-factor like activities through specific G-protein-coupled receptors (GPCR) on the cell surface [1]. Uncontrolled expression of this bioactive lipid or of its receptors has been linked to tumor growth and metastasis [1]. Stimulation with LPA culminates with the activation of several transcription factors, including NF-κB. Although the molecular basis for GPCR-mediated NF-κB activation continues to be elucidated, signal transduction was recently shown to share striking similarities with signaling trough antigen receptors in lymphocytes. In both cases, NF-κB signaling requires the activation of apical Protein Kinase C (PKC) and the subsequent assembly of a large signaling complex of CARMA1/3, BCL10 and MALT1 coined CBM complex [2–7]. The CBM complex serves as a docking platform for the recruitment and activation of the IκB kinase complex (IKK), which phosphorylates the NF-κB inhibitors IκBs, marking them for proteasomal degradation. NF-κB is then free to translocate into the nucleus and modulate the transcription of its target genes, driving lymphocyte activation and cytokines production [8].

In addition to its scaffolding function required for the activation of NF-κB, MALT1 is a paracaspase that cleaves a limited set of substrates to finely tune the immune response [9–12]. In the immune system, known MALT1 substrates count regulators of NF-κB, JNK/AP-1, mTOR, mRNA stability and of adhesion [8]. This enzymatic activity is crucial for the development of regulatory T cells, the differentiation of Th17 lymphocytes, and governs T-cell receptor (TCR)-driven proliferation and IL-2 production [9–11,13]. Interestingly, MALT1 protease activity was also found to modulate the integrity of the epithelial barrier in response to thrombin [3], therefore suggesting that MALT1 could be unleashed following GPCRs engagement.

Recently, we and others reported that the linear ubiquitin assembly chain complex (LUBAC) is dynamically recruited to the CBM following antigen receptor ligation and participates in IKK/NF-κB activation [14–17]. This triad composed of the E3 ligases HOIP (also known as RNF31), HOIL1 (also known as RCBK1) and the adaptor SHARPIN catalyzes linear ubiquitin chains and conveys NF-κB signaling downstream a variety of immunoreceptors [18]. In addition, HOIL1 was subsequently found to be cleaved and inactivated by MALT1 at the residue Arginine in position 165 [19–21]. Nevertheless, the contribution of the LUBAC to LPA-mediated NF-κB activation in non-immune cells has not been addressed. Here, by using embryonic fibroblasts derived from mice knockout for the LUBAC subunits, we unveiled a pivotal role for this complex during NF-κB activation in response to LPA. We also show that, paralleling the situation in lymphocytes, LPA triggers MALT1 protease activity and that the paracaspase cleaves HOIL1 at the Arginine 165. Although dispensable for IKK activation, this processing is required for the optimal production of the CBM target cytokine interleukin-6 (IL-6). Lastly, we report that the guanine exchange factor GEF-H1, which is activated by LPA [22], is required for MALT1 enzymatic activity and NF-κB activation in non-immune cells.

## Results and Discussion

The treatment of mouse embryonic fibroblasts (MEFs) with LPA, or with phorbol 12-myristate 13-acetate (PMA) plus ionomycin (PI) to mimic GPCR engagement, activates NF-κB in a CBM-dependent manner (Fig. 1A) [2,4,6,7]. To gain insights into the contribution of the LUBAC, we first used MEFs knockout for the catalytic subunit HOIP (*Tnf*^-/-^ *Hoip*^-/-^) and control *Tnf*^-/-^ *Hoip*^+/-^ cells [23]. Although LPA and PMA plus ionomycin led to a robust activation of NF-κB in *Tnf*^-/-^ *Hoip*^+/-^ cells, as demonstrated by the phosphorylation of the NF-κB inhibitor IκBα, this was not the case with *Tnf*^-/-^ *Hoip*^-/-^ cells, suggesting a defect in NF-κB activation (Fig. 1B). In line with this, the secretion of the CBM-dependent cytokine IL-6 was impaired without HOIP (Fig. 1C). Importantly, the genetic ablation of SHARPIN (chronic proliferative dermatitis, *Cpdm*, [24]) and HOIL1 (*Tnf*^-/-^ *Hoil1*^-/-^, [25]) also precluded the activation of NF-κB (Fig. 1D and 1E). Taken together, these data suggest that the LUBAC is required for NF-κB activation downstream of GPCR.

**Figure 1.**
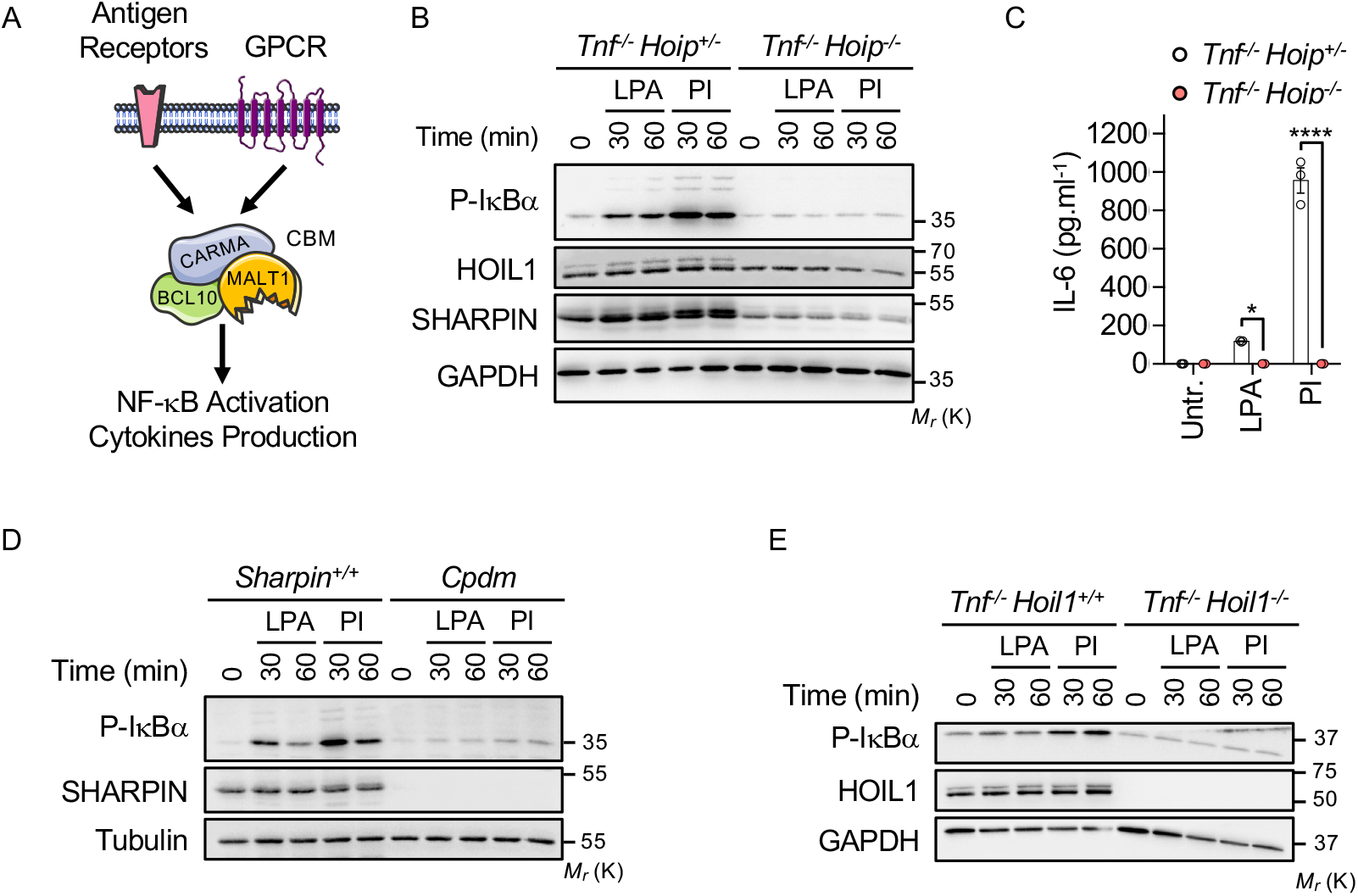
LPA-mediated NF-κB Activation requires the LUBAC. **A,** Graphical representation of NF-κB activation in response to antigen receptor engagement and to LPA stimulation. **B,** *Tnf*^-/-^ *Hoip*^-/-^ MEFs and their wild type counterparts *Tnf*^-/-^ *Hoip*^-/-^ were stimulated with 10 μg.mL^-1^ LPA or with 50 ng.mL^−1^ PMA plus 100 ng.mL^−1^ Ionomycin (PI), as indicated. Cell lysates were prepared and subjected to Western blotting analysis with antibodies specific for the indicated proteins. Molecular weight markers M_r_ (kDa) are shown. **C,** Abundance of IL-6, measured by ELISA in the supernatant of cells as in (B) stimulated for 16 hours (means ± SEM, n=3, ****P<0.0001, ANOVA). **D and E,** *Cpdm (D)*, *Tnf*^-/-^ *Hoil1*^-/-^ (E) and their wild type counterparts were treated as in (A). Lysates were prepared and subjected to Western blotting analysis as indicated. Data are representative of three independent experiments.

In lymphocytes, HOIL1 is rapidly cleaved by the paracaspase MALT1 after the arginine residue in position 165 [19–21]. Because MALT1 proteolytic activity was also proposed to be unleashed upon activation of the thrombin PAR1 GPCR in epithelial cells [3], we next assessed the status of HOIL1 in cells treated with LPA. We found that HOIL1 was cleaved in cells exposed to LPA or to PMA plus ionomycin (Fig. 2A). By contrast, stimulation with tumor necrosis factor-α (TNFα), a potent NF-κB activator that operates independently of MALT1, had no overt effect on HOIL1 (Fig. 2A). Consistent with a role for MALT1 enzyme, the inhibition of its catalytic activity with Mepazine [26,27] strongly reduced HOIL1 processing (Fig. 2B). GPCRs promote the activation of several signaling transduction pathways such as MAPK, Akt or NF-κB via the stimulation of guanine nucleotide exchange by heterotrimeric G proteins [28,29]. For instance, the ectopic expression of a constitutively active mutant of G_q_α (G_q_α_QL_) is sufficient to activate NF-κB in a BCL10-dependent manner [7]. We found that such mutant also drove the processing of FLAG-tagged HOIL1 when overexpressed in HEK293T cells (Fig. 2C). This was however not the case when the Arginine in position 165 on HOIL1 was substituted by a Glycine (HOIL1-R165G), hinting that the cleavage site of MALT1 is conserved upon GPCR engagement (Fig. 2C). Of note, similar results were obtained when the Kaposi Sarcoma-associated herpes virus (KSHV) GPCR (vGPCR), which encodes for constitutively active GPCR [30], was overexpressed. Altogether, our results suggest that MALT1 cleaves HOIL1 upon GPCR stimulation.

**Figure 2.**
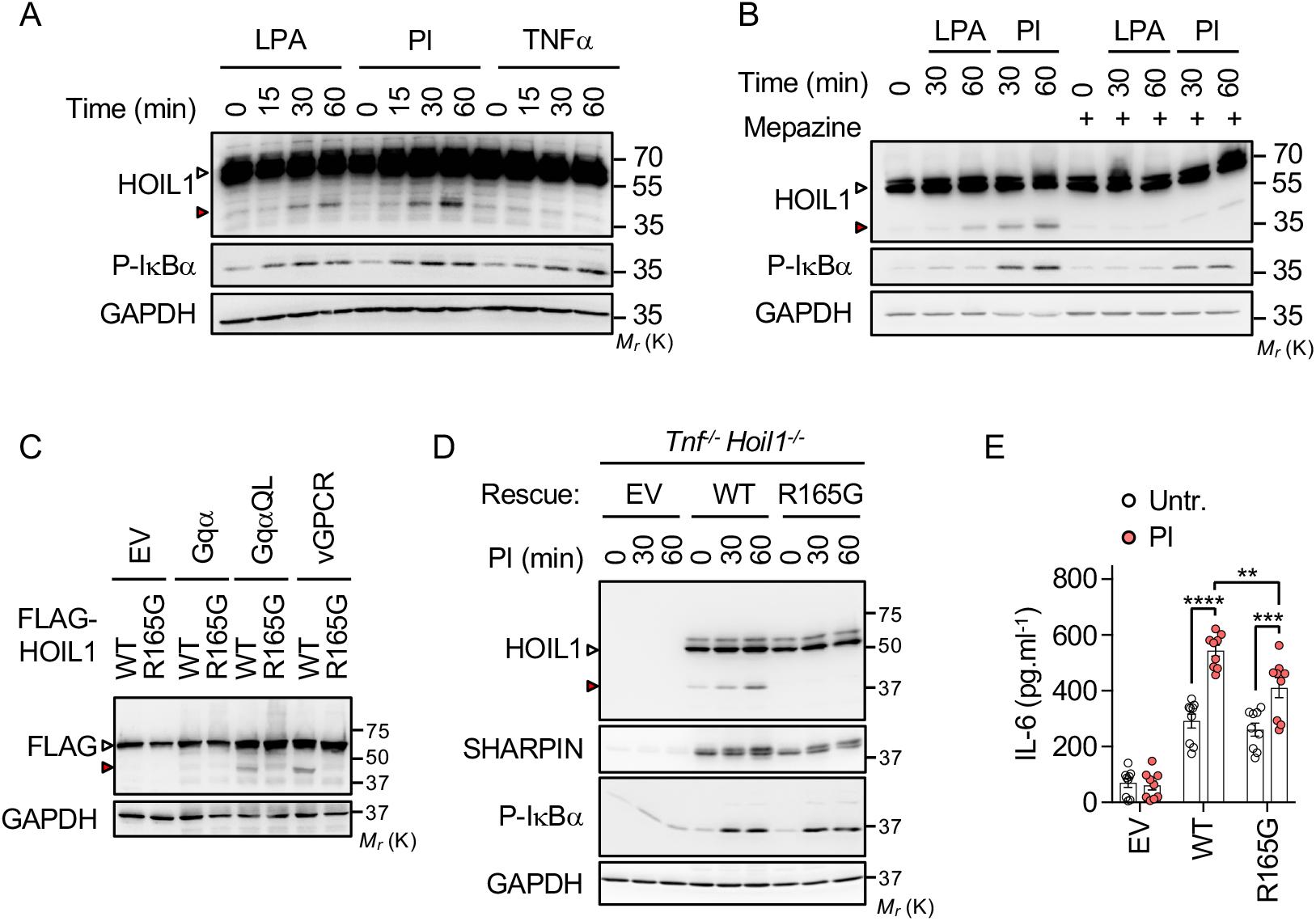
HOILl is cleaved by MALT1 following LPA Stimulation. **A,** MEFs were stimulated with 10 μg.mL^−1^ LPA, with 50 ng.mL^−1^ PMA plus 100 ng.mL^−1^ Ionomycin (PI), or with 10 ng.mL^−1^ TNFα as indicated. Lysates were subjected to Western blotting analysis with antibodies specific for the indicated proteins. White and red arrowheads show full-length and cleaved HOIL1, respectively. Molecular weight markers (M_r_) are shown. **B,** Western blotting analysis of MEFs pretreated with 20 μM of Mepazine for 90 minutes and stimulated as in (A). **C,** HEK293T cells were transfected with an empty vector (EV), or with plasmids encoding for G_q_α, G_q_α_QL_ or vGPCR together with FLAG-tagged HOIL1-WT or HOIL1-R165G. Lysates were prepared and Western blotting analysis was performed as indicated. **D,** *Tnf*^-/-^ *Hoil1*^-/-^ MEFs were retrovirally reconstituted with plasmids encoding for EV, HOIL1-WT or HOIL1-R165G, and stimulated as in (A). Cell lysates were prepared and subjected to Western blotting analysis, as indicated. **E,** Cells as in (D) were stimulated for 3 hrs. IL-6 secretion was analyzed by ELISA (means ± SEM. n=9, **P<0.01, ***P<0.001, ****P<0.0001, ANOVA). Data are representative of three independent experiments.

Because HOIL1 can negatively regulate antigen receptor-mediated NF-κB activation independently of IKK, unless cleaved by MALT1 [19], we next assessed the impact of HOIL1 cleavage in fibroblasts. To this end, MEFs *Tnf*^-/-^ *Hoil1*^-/-^ were retrovirally reconstituted with plasmids encoding for an empty vector (EV), HOIL1-WT or MALT1-resistant HOIL1-R165G (Fig. 2D). Both HOIL1-WT and HOIL1-R165G were expressed at levels comparable to parental cells, and restored SHARPIN levels supporting previous reports [31–33]. As expected, HOIL1 was cleaved upon PMA plus ionomycin treatment in HOIL1-WT, and not HOIL1-R165G expressing cells. Interestingly, the defect in IκBα phosphorylation observed without HOIL1 upon LPA or PMA plus ionomycin treatment was overcome in cells expressing HOIL1-WT or HOIL1-R165G, hinting at a normal activation of IKK (Fig. 2D). By contrast, HOIL1-R165G expressing cells did not secrete as much as IL-6 when compared to MEFs reconstituted with HOIL1-WT upon PMA plus ionomycin treatment (Fig. 2E), suggesting that HOIL1 processing is required for optimal IL-6 production. Together, our findings suggest a crucial role for the LUBAC in NF-κB activation downstream of GPCR. Echoing the situation in lymphocytes, MALT1 protease activity is unleashed and the paracaspase cleaves HOIL1 at the arginine 165. Although dispensable for IKK activation, HOIL1 processing appears required for the optimal secretion of the target IL-6.

LPA has been shown to drive the activation of the guanine exchange factor GEF-H1 (also called ARGHEF2) by releasing it from microtubules [22]. GEF-H1 exerts its varied functions through the activation of Rho GTPases. For instance, GEF-H1-driven activation of RhoA promotes actin polymerization, modulating cell contractility or endothelial barrier permeability [34–36]. Notably, GEF-H1 participates in the activation of NF-κB during intracellular pathogen recognition [37,38] or in response to microtubule destabilization [39]. Supporting previous works [37,38], we observed that the overexpression of GEF-H1 in HEK293T was sufficient to promote the transcriptional activity of NF-κB (Fig. 3A). Interestingly, we found that ectopic GEF-H1 also led to a robust cleavage of HOIL1 (Fig. 3B). Consistent with a contribution for GEF-H1 in MALT1 activity, MALT1-resistant HOIL1-R165G remained intact (Fig. 3B). In addition, MEFs deficient for GEF-H1 displayed reduced phosphorylation of IκBα and cleavage of HOIL1 in response to LPA and PMA plus ionomycin stimulations (Fig. 3C). As a consequence, the secretion of IL-6 was blunted (Fig. 3D). Collectively, these results suggest that GEF-H1 contributes to the CBM-driven activation of NF-κB following GPCR engagement.

**Figure 3.**
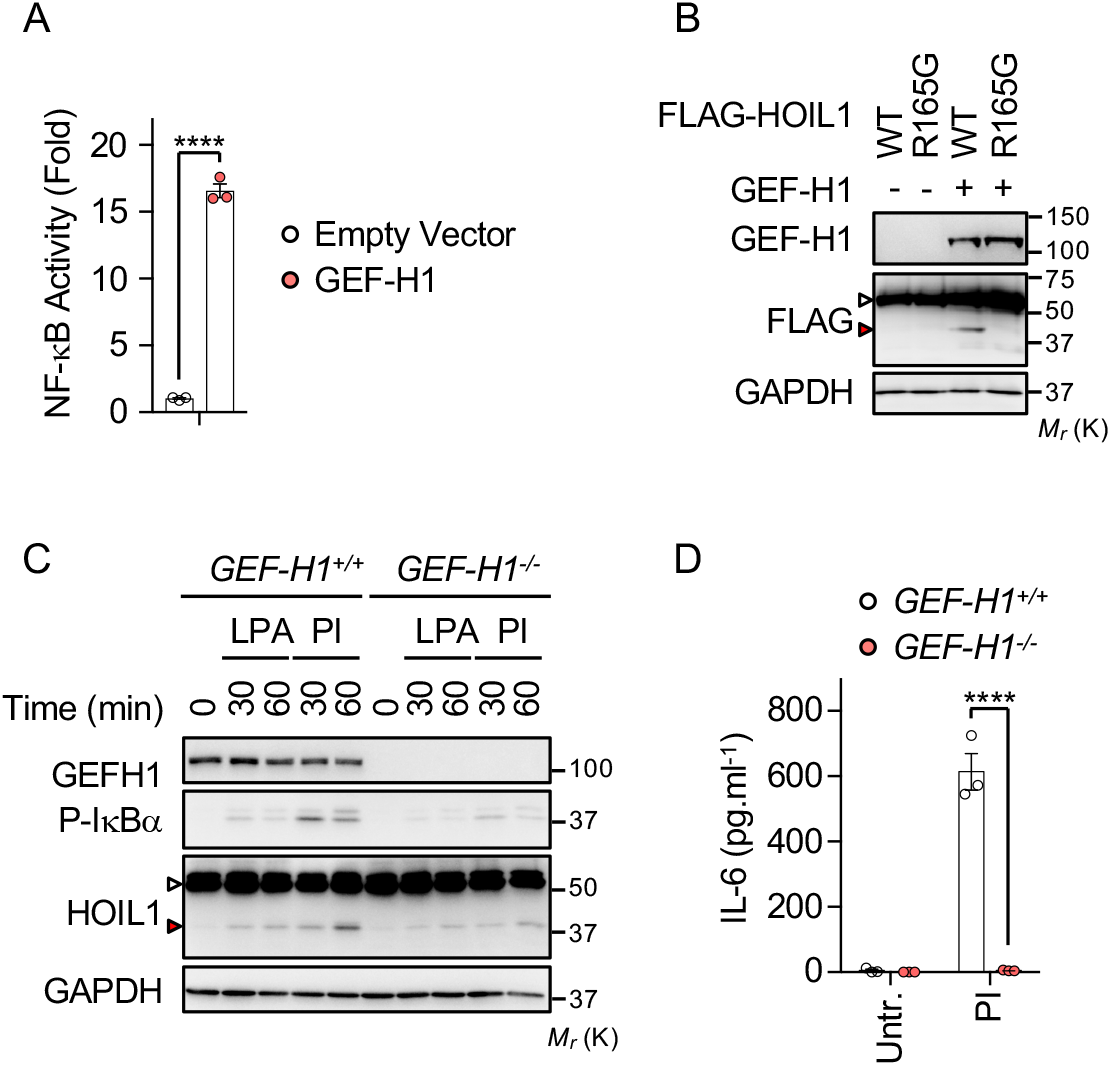
GEF-H1 participates in NF-κB activation and MALT1 Cleavage in response to LPA. **A,** HEK293T were transfected with a vector encoding for GEF-H1 or with an empty vector (EV) together with an NF-κB reporter plasmid. Shown is the mean ± SEM of triplicate experiments. ****P<0.0001 (ANOVA). **B,** HEK293T cells were transfected with an empty vector (EV), or with a GEF-H1 plasmid together with FLAG-tagged HOIL1-WT or HOIL1 - R165G. Cell lysates were subjected to Western blotting analysis with antibodies specific to the indicated proteins. White and red arrowheads show full-length and cleaved HOIL1, respectively. Molecular weight markers (M_r_) are shown. **C,** MEFs knockout for GEF-H1 (*GEF-H1*^-/-^) or control MEFs (*GEF-H1*^+/+^) were stimulated with 10 μg.mL^−1^ LPA or with 50 ng.mL^−1^ PMA plus 100 ng.mL^−1^ Ionomycin (PI) as indicated. Lysates were prepared and Western blotting analysis was performed as indicated. **D,** Abundance of IL-6, measured by ELISA in the supernatant of cells as in (C) stimulated for 3 hours (means ± SEM, n=3, ****P<0.0001, ANOVA). Data are representative of three independent experiments.

The CBM complex is an essential gateway for transducing NF-κB signaling downstream of antigen receptors and GPCRs [8]. In T and B lymphocytes, the CBM complex recruits the LUBAC to facilitate the optimal activation of NF-κB [14–17]. However, the situation in non-immune cells remains unknown. We now report that the LUBAC also contributes to LPA-induced NF-κB activation. This signaling node may therefore emerge as an appealing therapeutic target to jugulate GPCR signaling. The LUBAC has been reported to decorate key components of the antigen receptor-mediated NF-κB signaling pathway such as BCL10 and the IKK subunit NEMO with linear ubiquitin chains, and can also serve as an adaptor [14,16,17,40,41]. It will therefore be of clear interest to define the role of the catalytic activity of the LUBAC following GPCR engagement. Our results also unveil a dual role for HOIL1. Whereas HOIL1 positively conveys LPA-induced NF-κB activation, it lessens NF-κB signaling independently of IKK unless cleaved by the paracaspase MALT1. This is reminiscent of the situation in lymphocytes [19], and further investigation is now required to elucidate how exactly HOIL1 exerts its negative role. Lastly, we provide evidence that the guanine exchange factor GEF-H1 links LPA receptors to the CBM complex to promote the activation of MALT1 and NF-κB. This may provide a molecular framework to explain how the small GTPase RhoA, which can be activated by GEF-H1, drives NF-κB activation [42].

## Materials and Methods

### Cell culture and Reagents

HEK293T cells were purchased from the American Type Culture Collection (CRL-1573). MEFs deficient for HOIP (*Tnf*^-/-^ *Hoip*^-/-^) and HOIL1 (*Tnf*^-/-^ *Hoil1*^-/-^) as well as their wild-type counterparts were kindly provided by H. Walczak (University College London, London, United Kingdom) [23,25]. MEFs deficient for SHARPIN (chronic proliferative dermatitis, *Cpdm*) and their wild-type counterparts were a generous gift by J. Ivaska (University of Turku, Turku, Finland) [24]. MEFs deficient for GEFH1 and their wild-type littermates were previously described [22,43]. All cell lines were cultured in Dulbecco’s modified Eagle’s media (DMEM, Life Technologies) supplemented with 10% FBS and Penicillin/Streptomycin (Life Technologies). Cells were stimulated with 10 μg.mL^−1^ of LPA (Santa Cruz), with a mixture of 50 ng.mL^−1^ PMA (Sigma) and 100 ng.mL^−1^ Ionomycin (Sigma), or with 10 ng.mL^−1^ TNFα (R&D Systems). The protease activity of MALT1 was blocked with 20 μM of Mepazine (Chembridge). Mouse IL-6 was detected from cell culture supernatants by ELISA according to the manufacturer’s instructions (R&D Systems).

### Western Blotting Analysis and Antibodies

Cells were washed with ice-cold PBS and lysed with TNT buffer [50 mM Tris-HCl (pH 7.4), 150 mM NaCl, 1% Triton X-100, 1% Igepal, 2 mM EDTA] supplemented with protease inhibitors (Thermo Fisher Scientific) for 30 min on ice. Samples were cleared by centrifugation at 9,000g and protein concentration determined by a BCA assay (Thermo Fisher Scientific). 5-10 μg denaturated proteins were resolved by SDS-PAGE and transferred to nitrocellulose membranes (GE Healthcare). Antibodies against HOIL1 (H-1, 1:1,000), GAPDH (6C5, 1:20,000) were from Santa Cruz. Antibodies to phosphorylated IκBα (5A5, 1:5,000), and GEF-H1 (55B6, 1:1,000) were from Cell Signaling Technology. Antibodies against SHARPIN (A303-559A, 1:5,000) from Bethyl Laboratories and against FLAG (M2, 1:5,000) from Sigma were also used. Horseradish peroxidase (HRP)-conjugated secondary antibodies were purchased from Southern Biotechnology.

### Expression Plasmids and Transfections

Plasmids encoding for G_q_α, its constitutively active mutant G_q_α_QL_ and vGPCR were a gift from J.S. Gutkind [44]. HOIL1-WT and HOIL1-R165G in a pCDH1-MSCV-EF1α-GreenPuro vector were previously described [19]. GEF-H1 (pCMV-ARHGEF2, Origene) was cloned into a pCDH1-MSCV-EF1α-GreenPuro plasmid (SBI). HEK293T cells were transfected using a standard calcium phosphate protocol, as previously described [45,46]. NF-κB luciferase gene reporter assays were performed according to the manufacturer’s instruction (Promega), as previously described [19]. For infections, HEK293T were transfected with plasmids encoding for HOIL1-WT, HOIL1-R165G or an empty vector together with pVSV-G and psPAX2 plasmids. Cell culture supernatants containing lentiviral particles were collected after 48 hours and added on *Tnf*^-/-^ *Hoil1*^-/-^ MEFs with 8 μg.mL^−1^ polybrene (Santa Cruz) during a 2,500 x *rpm* centrifugation for 90 minutes. 24 hours postinfection, cells were selected with 1 medium containing 1 μg.mL^−1^ of puromycin.

### Statistical Analysis

Statistical analyses were performed using GraphPad Prism 7 software using two-way ANOVA and *P* values are indicated in the figure legends.

## Acknowledgments

We thank H. Walczak, J. Ivaska and J.S. Gutkind for kindly providing reagents. This research was funded by an International Program for Scientific Cooperation (PICS, CNRS), Fondation pour la Recherche Médicale (Equipe labellisée DEQ20180339184), Fondation ARC contre le Cancer (NB), Ligue nationale contre le cancer comités de Loire-Atlantique, Maine et Loire, Vendée (JG, NB), Région Pays de la Loire et Nantes Métropole under Connect Talent Grant (JG), the National Research Agency under the Programme d’Investissement d’Avenir (ANR-16-IDEX-0007), and the SIRIC ILIAD (INCa-DGOS-Inserm_12558). TD is a PhD fellow funded by Nantes Métropole.

## Author Contributions

Conceptualization, TD and NB; Methodology, TD and NB; Investigation, TD, SC and NB; Resources, RR; Supervision, JG; Writing-Original Draft, TD and NB; Writing-Review & Editing, TD, JG, and NB; Funding Acquisition, JG and NB.

